# Patterns and mechanisms of diminishing returns from beneficial mutations

**DOI:** 10.1101/467944

**Authors:** Xinzhu Wei, Jianzhi Zhang

## Abstract

Diminishing returns epistasis causes the benefit of the same advantageous mutation smaller in fitter genotypes, and is frequently observed in experimental evolution. However, its occurrence in other contexts, environment-dependence, and mechanistic basis are unclear. Here we address these questions using 1005 sequenced segregants generated from a yeast cross. Under each of 47 examined environments, 66-92% of tested polymorphisms exhibit diminishing returns epistasis. Surprisingly, improving environment quality also reduces the benefits of advantageous mutations even when fitness is controlled for, indicating the necessity to revise the global epistasis hypothesis. We propose that diminishing returns originates from the modular organization of life where the contribution of each functional module to fitness is determined jointly by the genotype and environment and has an upper limit, and demonstrate that our model predictions match empirical observations. These findings broaden the concept of diminishing returns epistasis, reveal its generality and potential cause, and have important evolutionary implications.

## INTRODUCTION

Diminishing returns epistasis refers to a reduction in the benefit of an advantageous mutation when it occurs in a relatively fit genotype compared with that in a relatively unfit genotype (Griffing 1950; Jerison and Desai 2015). It is believed to explain at least in part why experimental evolution of microbes almost invariantly shows a decreasing speed of adaptation as the fitness of the population rises (Wiser, et al. 2013; Couce and Tenaillon 2015). Diminishing returns epistasis has been indirectly inferred from the dynamics of adaptation (Moore, et al. 2000; Kryazhimskiy, et al. 2009; Perfeito, et al. 2014; Good and Desai 2015) and directly demonstrated by engineering the same mutation in multiple strains of different fitnesses (MacLean, et al. 2010; Chou, et al. 2011; Khan, et al. 2011; Kryazhimskiy, et al. 2014; Wang, et al. 2016). While diminishing returns epistasis appears common among fixed mutations in experimental evolution, it is unknown whether it is restricted to experimental evolution, where fixed beneficial mutations are *de novo* and tend to have large effects (Orr 2002; Rokyta, et al. 2005), or is also widespread among standing genetic variants. Furthermore, whether and how the pattern of diminishing returns epistasis varies across environments have not been investigated. Most importantly, the underlying cause of diminishing returns epistasis remains elusive. A commonly considered hypothesis termed the global epistasis hypothesis posits that “the effect of each mutation depends on all other mutations, but only through their combined effect on fitness” and that “each individual beneficial mutation provides a smaller advantage in a fitter genetic background” (Kryazhimskiy, et al. 2014). Although this hypothesis is currently regarded as the leading description and explanation of diminishing returns epistasis (Kryazhimskiy, et al. 2014; Wang, et al. 2016), to what extent it is true and why it may be true remain unanswered. Note that the diminishing returns relationship between the activity of an enzyme and the flux of the relevant metabolic pathway is well explained by the metabolic control theory (Kacser and Burns 1981; Dykhuizen, et al. 1987; Chou, et al. 2014), but this theory cannot explain diminishing returns epistasis arising from interactions among mutations of different genes.

Here we develop a high-throughput method to investigate diminishing returns epistasis among standing genetic variants. We report widespread diminishing returns epistasis from single nucleotide polymorphisms (SNPs) segregating in budding yeast, discover a novel type of diminishing returns that results from an improvement in environment quality, provide evidence that the origin and patterns of diminishing returns are best explained by the modular structure of life, and discuss evolutionary implications of these findings.

## RESULTS

### Quantifying diminishing returns epistasis by comparing mean benefits in multiple genetic backgrounds

Diminishing returns epistasis is conventionally demonstrated by showing that the same mutation causes a smaller growth rate increase in a relatively fit strain than in a relatively unfit strain (MacLean, et al. 2010; Chou, et al. 2011; Khan, et al. 2011; Kryazhimskiy, et al. 2014; Wang, et al. 2016). If the observed diminishing returns epistasis is genuine and general, it should also be testable by comparing the mean benefits of the mutation in two sets of strains that differ in mean growth rate (see Materials and Methods). Using this approach allows testing diminishing returns epistasis for each nucleotide difference between the genomes of two organisms that can be crossed to produce a hybrid and its segregants, as long as the genotypes and growth rates of the segregants can be acquired. For example, for an A/G polymorphism at a site, we can calculate the effect of substituting A with G by comparing the mean growth rate of segregants with genotype A (or AA for diploid segregants) and the mean growth rate of segregants with genotype G (or GG for diploid segregants) at the site, because the A segregants and G segregants are on average equivalent for the rest of their genomes due to random assortment and recombination in meiosis. The above calculation can be separately performed in two sets of strains with different mean growth rates, allowing testing diminishing returns epistasis.

We applied this method to a dataset that includes the genome sequences of 1005 haploid segregants produced from the hybrid between the BY and RM strains of the yeast *Saccharomyces cerevisiae* (Bloom, et al. 2013). BY is derived from the widely used laboratory strain S288c, whereas RM is derived from the vineyard strain RM11-1a. The dataset also includes the mean end-point colony radius of each segregant on agar plates in 47 environments, which vary in temperature, pH, carbon source, metal ions, and small molecules (Bloom, et al. 2013). We estimated the growth rate of a segregant in each environment using the corresponding colony radius (**Fig. S1**; see Materials and Methods).

### Widespread diminishing returns epistasis among standing genetic variants

To demonstrate diminishing returns epistasis, we need to show that the mean benefit of a mutation in slow-growth segregants is greater than that in fast-growth segregants. In each environment, we computed for each SNP the mean growth rate (*R*_BY_) of the least fit 20% of BYallele-carrying segregants and that (*R*_RM_) of the least fit 20% of RM-allele-carrying segregants (**Fig. 1a**). The effect of the SNP in these slow-growth segregants is *s*_L_ = |*R*_BY_ - *R*_RM_|. We similarly computed the mean growth rate (*R*′_BY_) of the fittest 20% of BY-allele-carrying segregants and that (*R*′_RM_) of the fittest 20% of RM-allele-carrying segregants (**Fig. 1a**) and estimated the effect of the same SNP in these fast-growth segregants by *s*_H_ = |*R’*_BY_ - *R’*_RM_|. The diminishing returns epistasis is present if *s*_H_ < *s*_L_.

**Fig. 1.**
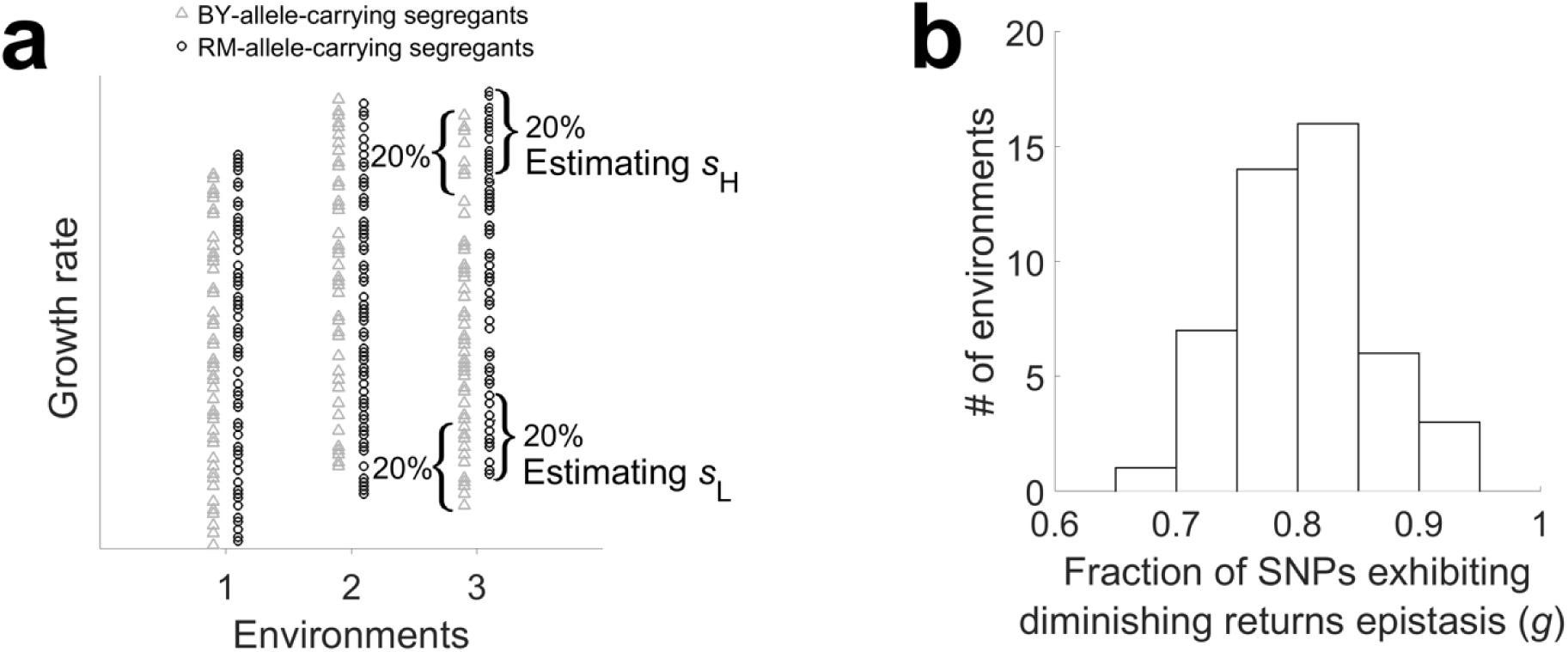
Widespread diminishing returns among standing genetic variants in yeast. *s*_H_, growth rate effect of a SNP in fast-growth segregants; *s*_L_, growth rate effect of a SNP in slow-growth segregants. (**a**) Scheme for estimating *s*_H_ and *s*_L_. For each SNP under each environment, grey triangles represent BY-allele-carrying segregants, while black circles represent RM-allele-carrying segregants. The fittest 20% of BY-allele-carrying and fittest 20% of RM-allele-carrying segregants are used to estimate *s*_H_, whereas the least fit 20% of BY-allele-carrying and least fit 20% of RM-allele-carrying segregants are used to estimate *s*_L_. The data plotted are hypothetical and not all segregants used in each group are shown. (**b**) Frequency distribution of the fraction (*g*) of SNPs exhibiting diminishing returns epistasis (i.e., *s*_H_ < *s*_L_) in the 47 environments examined.

Under the null hypothesis that the benefit of a mutation is independent of the growth rate of the genetic background, a SNP has a 50% chance to exhibit *s*_H_ < *s*_L_. Strikingly, in each of the 47 environments studied, between *g* = 66% and 92% of the 28,220 SNPs tested show *s*_H_ < *s*_L_(**Fig. 1b**, **Table S1**), with an average of 80%. Although the relationship between *s*_H_ and *s*_L_ for one SNP is not independent from that for a linked SNP, the estimated *g* in each environment is unbiased. The non-independence among SNPs, however, makes it difficult to test if *g* significantly exceeds the chance expectation of 50% in an environment. But, because the growth rates of all segregants were separately measured in different environments, the *g* values from different environments were estimated independently. The observation that all 47 independently estimated *g* values exceed 50% has a binomial probability lower than 10^-14^ under the null hypothesis of *g* = 0.5, strongly suggesting a general presence of diminishing returns epistasis across environments. Our results hold when 10% or 30% (instead of 20%) of segregants are used to estimate *s*_L_ and *s*_H_. We verified that our results are not an artifact of transforming colony radius to growth rate, because repeating the analysis using colony radius showed that *g* varies from 58% to 88% across the 47 environments. We also examined in each environment the subset of SNPs for which the beneficial allele is the same in the 20% slow- and 20% fast-growth segregants. Forty of the 47 environments (*P* = 10^-6^, binomial test) have more than 50% of such SNPs exhibiting diminishing turns (mean across 47 environments = 62%).

Following a recent analysis of the same dataset (Wei and Zhang 2017), we mapped quantitative trait loci (QTLs) underlying the growth rate variation among the segregants (at a false discover rate of 0.05) in each of the 47 environments. The number of QTLs identified ranged from 0 to 33 in the 47 environments, with a mean of 15.8 (Wei and Zhang 2017). Zero QTL was mapped in one environment. In three environments, the ratios between the number of BY-allele-carrying segregants with growth data and that of RM-allele-carrying segregants with growth data at putative QTLs differ significantly from the corresponding ratio in all 1005 segregants, making effect size estimation unreliable. In two environments, *s*_H_ < *s*_L_ in exactly 50% of QTLs. In the 41 remaining environments, 37 showed *s*_H_ < *s*_L_ in over 50% of QTLs (*P* = 5.2×^-8^, binomial test) and 20 of them showed *s*_H_ < *s*_L_ in significantly more than 50% of QTLs (nominal *P* < 0.05; **Table S1**). By contrast, only 4 environments showed *s*_H_ < *s*_L_ in fewer than 50% of QTLs and none of these environments showed *s*_H_ < *s*_L_ in significantly fewer than 50% of QTLs. Thus, the prevalence of diminishing returns epistasis is also evident among SNPs having independent growth effects. By bootstrapping the segregants used (see Materials and Methods), we confirmed that *s*_H_ is significantly smaller than *s*_L_ at the nominal *P*-value of 0.05 for 202 of a total of 613 QTLs with unbiased growth rate distributions (101 significant QTLs after Bonferroni correction of multiple testing per environment).

### Benefits of advantageous mutations decrease with environment quality

Because the yeast growth data contained 47 environments, we investigated how environmental changes impact the growth effects of advantageous mutations. Specifically, we wonder if improving the environment also leads to the diminishing returns phenomenon. To this end, we define the quality (*Q*) of an environment to the population of segregants considered by the mean growth rate of all segregants in the environment (**Table S1**). We first measured the effect (*s* > 0) of each SNP in an environment by the absolute value of the difference between the mean growth rate of all BY-allele-carrying segregants and that of all RM-allele-carrying segregants in the environment. If having better environments reduces the benefit of an advantageous mutation, *s* should decrease as *Q* rises (see **Fig. 2a** for an example). Such a negative correlation between *Q* and *s* should be common among all SNPs examined if this type of diminishing returns is widespread. Indeed, for 92.7% of SNPs across the genome, we observed a negative rank correlation between *Q* and *s* across environments (**Fig. 2b**).

**Fig. 2.**
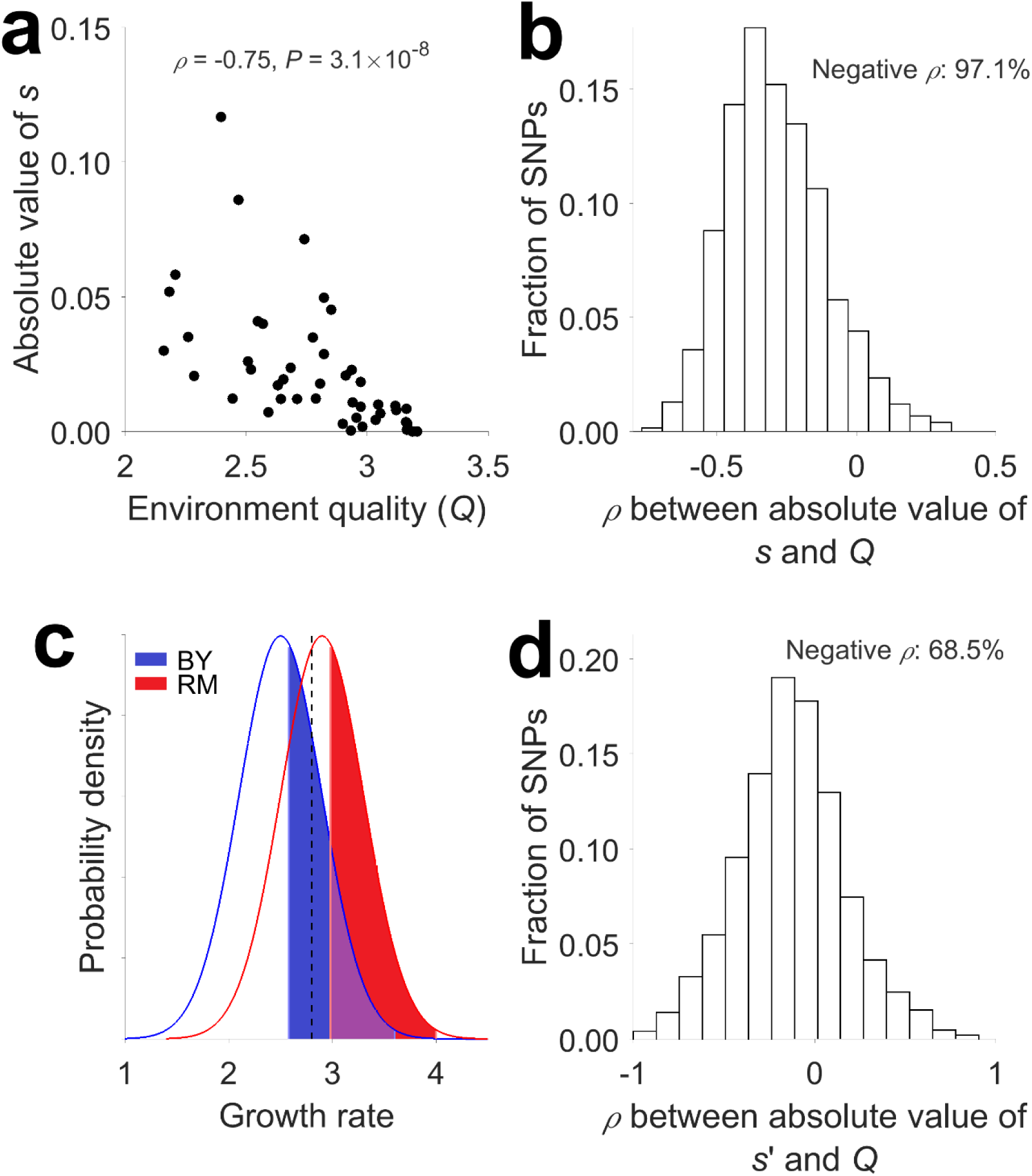
Most SNPs *s*_H_ ow a negative correlation between its effect on growth rate and environment quality (*Q*). (**a**) Among-environment correlation between *Q* and the absolute value of the growth effect for the SNP exhibiting the strongest negative correlation. Each dot represents one environment. Spearman’s rank correlation *ρ* and the associated *P*-value are presented. (**b**) Frequency distribution of the rank correlation (*ρ*) between *Q* and the absolute value of the growth effect (*S*) of a SNP measured using all segregants. (**c**) Scheme for estimating the growth effect (*s’*) of a SNP upon the control of growth rate. After the focal growth rate (vertical dashed line) is chosen, the same percentile ranges (blue shaded area for BY and red shaded area for RM) are used in the growth rate distributions of BY- and RM-allele-carrying segregants for estimating the effect size. See main text for details. (**d**) Frequency distribution of the rank correlation between *Q* and the absolute value of the growth effect (*s’*) of a SNP upon the control of growth rate.

To examine whether the above negative correlation is simply a byproduct of the canonical diminishing returns epistasis associated with a rise in the growth rate of the background genotype, we controlled for growth rate in estimating *s* using the following procedure (**Fig. 2c**). Firstly, we chose a fixed growth rate *R* for all environments. Secondly, we picked two groups of 20% of all BY-allele-carrying segregants for each SNP and in each environment. The first group are the least fit segregants whose growth rates exceed *R*, while the second group are the fittest segregants whose growth rates are below *R*. Thirdly, we estimated the mean growth rate *R*_BY_ using these 40% of BY-allele-carrying segregants and identified the percentile range of these segregants among all BY-allele-carrying segregants. Fourthly, we estimated the corresponding *R*_RM_ from the segregants in the same percentile range of all RM-allele-carrying segregants. Fifthly, we calculated the fitness effect of the SNP by *s*_1_ = |*R*_BY_ – *R*_RM_|. Sixthly, we repeated the above steps 2-5 after switching BY with RM, and estimated *R’*_RM_, *R’*_BY_, and *s*^2^ = |*R’*_RM_ - *R’*_BY_|. We then estimated *s’* = (*s*_1_ + *s*_2_)/2, which is the growth effect of the SNP in the environment given *R*. In this procedure, the same percentile range in BY- and RM-allele-carrying segregants were used to ensure an unbiased estimation of the effect size of each SNP upon the control of growth rate.

Because the range of growth rates of all segregants varies substantially among the 47 environments (**Fig. S1**), to investigate the correlation between *Q* and *s’* among as many environments as possible, we picked an *R* of 2.8, which allowed a comparison among nine environments. We found that the rank correlation between *Q* and *s’* is negative for 68.5% of all SNPs examined (**Fig. 2d**). We repeated this analysis under two other *R* values (2.6 and 3.0), and found the average fraction of SNPs exhibiting negative correlations with *Q* to be 63% for the three *R* values considered. A total of 600 SNPs were identified as QTLs in one or more environments. For each of these SNPs, we estimated its effect (*s* and *s’*) in each environment and correlated it with *Q*. For the 600 correlations between *s* (or *s’*) and *Q* across the 47 environments, 86.2 % (or 72.3%) are negative. These results suggest that, although variable fitness is a cause of the negative *Q-s* correlation, it is not the sole reason. Hence, the diminishing returns from advantageous mutations in better environments, a form of gene-environment interaction (G×E), is distinct from the canonical diminishing returns in fitter genotypes within an environment, a form of gene-gene interaction (G×G).

Although the above results from SNPs and those from QTLs are qualitatively similar, they differ quantitatively. We confirmed that SNP density does not affect the fraction of SNPs showing negative *Q-s* correlations, nor does it affect the average *Q-s* correlation (see Materials and Methods). We discuss factors that may cause the quantitative differences between SNP- and QTL-based results in Discussion.

### The modular life model recapitulates the empirical patterns of diminishing returns

The above finding that the same mutation confers different advantages on different genetic backgrounds even when these backgrounds are equally fit cannot be explained by the global epistasis hypothesis and suggests the relevance of the specific genomic compositions of these backgrounds to the fitness effect of a mutation. It is widely accepted that life is organized in a highly modular manner, where each module is a discrete object composed of a group of tightly linked components and performs a relatively independent task (Raff 1996; Hartwell, et al. 1999; Ihmels, et al. 2002; Ravasz, et al. 2002; Barabasi and Oltvai 2004; Wall, et al. 2004; Wagner, et al. 2007). Intuitively, diminishing returns epistasis could arise from the modular structure of life. Specifically, our modular life model posits that each module makes a distinct contribution to fitness and that this contribution has an upper limit. Under this model, the same advantageous mutation may contribute to a module and fitness greatly if the functionality of the module is far from its maximum but may contribute only slightly if the module is approaching its maximal functionality. In addition, we assume that the environment contributes differently to the functionalities of various modules and that different environments have different contributions. Because the functionalities of various modules can be different among equally-fit genotypes, under this model, the specific genomic composition of the background genotype matters to the fitness effect of a mutation. Our model can be seen as an extension of the global epistasis model from one module to multiple modules and from considering genetic effect only to considering both genetic and environmental effects. This extension presumably allows making specific testable predictions about diminishing returns epistasis both within and across environments. We here explore the modular life model in an attempt to recapitulate the major empirical patterns of diminishing returns.

We started by a computer simulation of the modular life model (**Fig. 3a** and Materials and Methods). We considered two scenarios where the growth rate of a genotype is respectively determined by the geometric mean functionality of all modules or the arithmetic mean functionality of all modules. The results obtained under the two scenarios are qualitatively similar, and they are respectively presented in the main text (**Fig. 3**) and **Fig. S2**. According to the modular life model, we simulated the genotypes and growth rates of 1000 haploid segregants in 50 environments (see Materials and Methods). One hundred genes belonging to 10 modules were considered, with each gene harboring one SNP that distinguishes between a fully functional allele and a null allele. We analyzed the simulated data the same way we analyzed the real data. Similar to what was observed in the real data (**Fig. 1b**), the simulated data show diminishing returns epistasis for >50% of SNPs in each environment (**Fig. 3b**). Furthermore, similar to what was found in the real data (**Fig. 2b, d**), most SNPs in the simulated data show a negative correlation between growth effect and environment quality, with or without the control for growth rate (**Fig. 3c**). The similarity between the results from the simulated data and real data indicates that the observed patterns of diminishing returns are explainable by the modular feature of life.

**Fig. 3.**
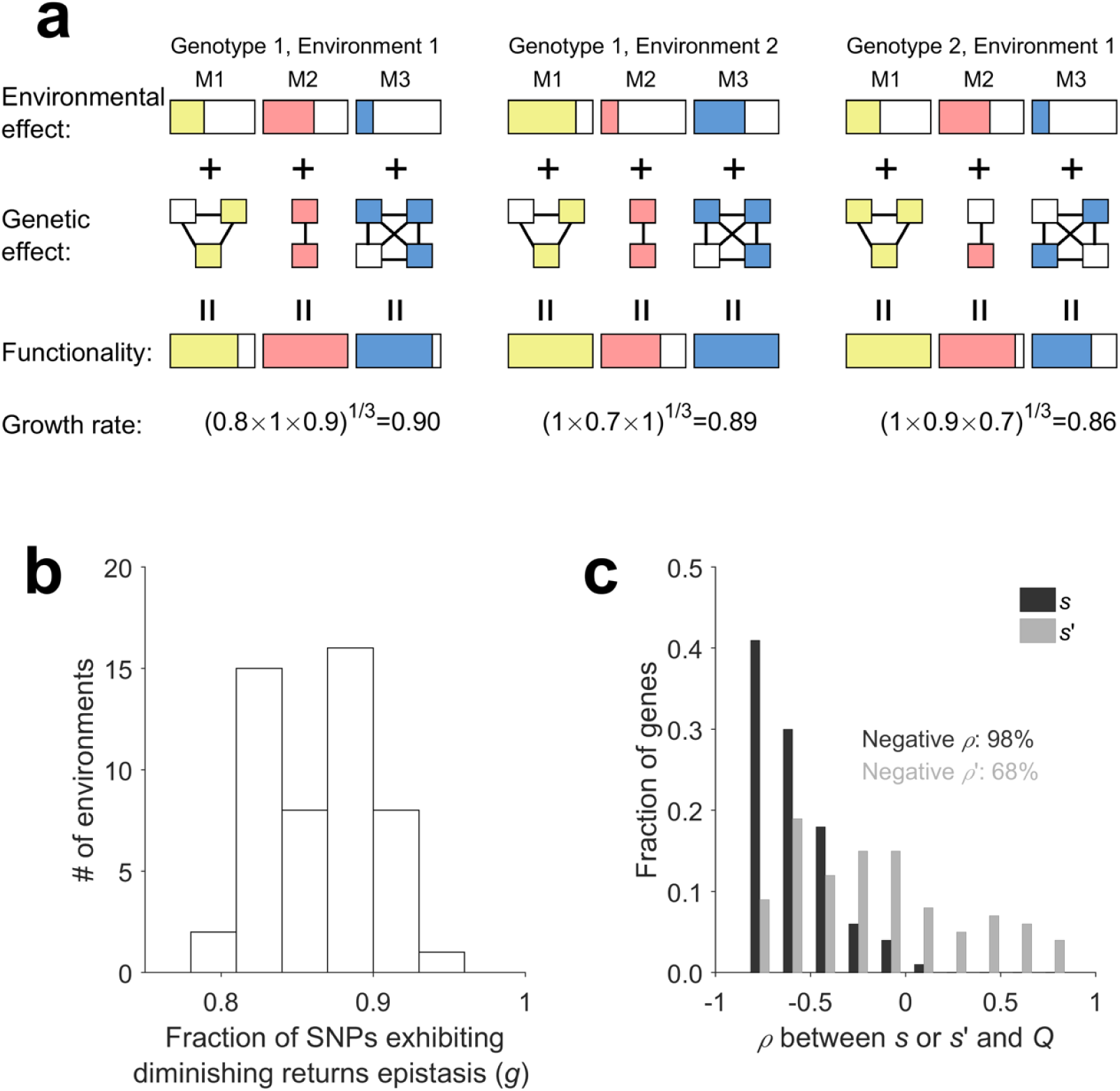
Simulation of the modular life model produces diminishing returns patterns resembling empirical observations. (**a**) Simulation scheme under the geometric mean growth rate model. Different modules (M1, M2, and M3) are colored differently. Different environments (Environments 1 and 2) contribute differently to various modules, as illustrated by the three boxes that are filled to different levels. Each module contains a number of genes, each of which could have either a functional allele designated as 1 (filled box) or a null allele designated as 0 (open box). Two genotypes (Genotypes 1 and 2) are shown as examples. The functionality of a module is the sum of environmental and genetic contributions but cannot exceed 1. The growth rate of each genotype is computed from the functionalities of the individual modules using the formula indicated. See Methods for parameters used in the simulation. (**b**) Frequency distribution of the fraction of genes exhibiting diminishing returns from simulated data. (**c**) Frequency distribution of the rank correlation (*ρ*) between *Q* and the effect of a SNP measured using either all segregants (*s*; black) or a group of segregants with a fixed median growth rate (*s’*; grey). The fraction of *ρ*’s that are negative is indicated in black and grey for s and **s’**, respectively. Here, *s* and *s’* could be negative if the functional allele is found less fit than the null allele (due to sampling error).

### Why effect size decreases with environment quality even after the control for growth rate

Although the canonical diminishing returns epistasis is easily explained by the modular life model, that *s’* decreases with *Q* (**Fig. 2d** and **Fig. 3c**) is puzzling. Furthermore, because we estimated *s’* from groups of segregants that differ in multiple genes, it is unclear whether the negative correlation between *s’* and *Q* holds when *s’* is estimated by comparing genotypes that differ by a single SNP upon the control of growth rate across environments. To this end, we measured the effect of a beneficial mutation in each genetic background and then averaged this effect across multiple backgrounds in simulated data. Specifically, we simulated 50,000 segregants in 50 environments as in the previous section except that stochastic noise in growth rate is omitted to improve the sensitivity of the analysis. In each environment, we first identified all segregants whose growth rates are in the range of 0.899-0.901. This range is much narrower than the maximal growth effect of any beneficial mutation simulated; therefore, the identified segregants are essentially equally fit. We estimated the growth effect of a gene in an environment by averaging the effect of replacing its null allele with functional allele in the above segregants in which the focal gene is occupied by the null allele. We then correlated between the growth effect of the gene and *Q* among environments. For the 100 genes simulated, 63 showed a negative rank correlation (*P* = 0.006, *N* = 100, binomial test). We repeated this analysis using another growth rate range (0.949-0.951) and found 68 of 100 genes to show negative rank correlations (*P* = 2×10^-4^). These results, quantitatively similar to those in **Fig. 3c**, confirm that the negative correlation between *s’* and *Q* observed in the simulation is genuine. The cause for this correlation is that, when the growth rate is controlled for, the among-module variance in functionality increases with *Q*. The reason is that, in this scenario, under a high *Q* environment, genotype quality must be relatively low, meaning that it has only a small number of functional alleles distributed among all modules, rendering the among-module variance in functionality relatively high. By contrast, under a low *Q* environment, genotype quality must be relatively high, meaning that it has many functional alleles distributed among all modules, rendering the among-module variance in functionality relatively low. Thus, the fraction of modules approaching the upper limit in functionality is greater in good environments than in poor environments, even when the mean functionality per module is the same. Consequently, the growth effect of a beneficial mutation tends to reduce with *Q*. We confirmed this reasoning using the above simulated data. In particular, we found that the among-module variance in functionality averaged across all segregants aforementioned correlates positively with *Q* for both of the growth rate ranges considered (**Fig. S3a**, **b**). The same trend holds when growth rate is defined by the arithmetic mean instead of geometric mean of functionality across modules (**Fig. S3c**, **d**).

## DISCUSSION

In this work, we designed a high-throughput method for testing diminishing returns epistasis among standing genetic variants and applied it to 28,220 SNPs as well as 741 QTLs between two yeast strains. We found widespread diminishing returns from beneficial mutations in each of the 47 environments studied, demonstrating that diminishing returns epistasis is abundant among natural genetic variants. There are pros and cons in analyzing QTLs only versus analyzing all SNPs. The QTL-based analysis considers influential SNPs that are independent from one another, but undoubtedly misses many causal SNPs due to the limited statistical power in QTL identification and hence provides an incomplete picture of the entire genome. The analysis of all SNPs provides a complete and unbiased picture of the genome, but because of the linkage among SNPs, some of the statistical tests are difficult. Nevertheless, we found that the two approaches resulted in qualitatively similar findings. The minor quantitative differences between the two approaches could be due to chance and/or the following. First, QTLs tend to have much larger effects than average SNPs. According to the modular life model, larger effects are more likely to occur in modules that are still far from saturation in functionality. Thus, QTLs are less likely than average SNPs to be subject to diminishing returns epistasis. Second, the commonly used QTL mapping assumes an additive model, which may underestimate the diminishing returns. If QTLs that show diminishing returns within an environment also tend to have diminishing returns when *Q* rises, the *Q-s* correlation will also be affected.

Canonical diminishing returns epistasis is a form of gene-gene interaction, because it is conventionally quantified by comparing the effect of a mutation in genotypes of different fitnesses in the same environment. Our work broadens the concept of diminishing returns to gene-environment interaction, because we found that the effect of a beneficial mutation decreases with environment quality. The results suggest that both types of diminishing returns (gene-gene and gene-environment interactions) are prevalent among standing genetic variants across environments. Our observation supports the common belief that the fitness effects of mutations tend to increase in stressful environments (Agrawal and Whitlock 2010) and further demonstrates that this increase also occurs even when the background genotype fitness is controlled.

The prevailing view prior to this study is that diminishing returns depends on the fitness of the background genotype, as described by the global epistasis hypothesis. Our finding that the benefit of an advantageous mutation decreases with environment quality even when the fitness of the background genotype remains unchanged cannot be explained by the current global epistasis model. Furthermore, a close examination of a previous study (Kryazhimskiy, et al. 2014) showed that the growth effects of a mutation in several strains of similar growth rates can be significantly different even under the same environment (**Table S2**). We proposed that diminishing returns can be explained by the modular structure of life, where each module contributes to a fitness component and has a maximal possible contribution. Consistently, our computer simulation demonstrates that this modular life model recapitulates the empirical patterns of diminishing returns. The patterns of diminishing returns observed in the simulation are due to three properties of the model that are not dependent on the specific parameters used in the simulation. First, when fitness is defined by the geometric mean of functionality among modules, the geometric mean generates a concave relationship between the functionality of a module and fitness. Second, when the functionality of a module saturates, any mutation that could potentially increase the functionality of the module becomes neutral. This is why diminishing returns epistasis is still observed even when fitness is defined by the arithmetic mean functionality across modules (**Fig. S2**). Third, because better environments on average have higher environmental contributions to the functionality of a module, the functionality of a module is closer to saturation in better environments. Consequently, the second property dictates that the benefit of an advantageous mutation is expected to be smaller in better environments. Fourth, as explained in the last section in RESULTS, even when fitness is controlled for, the variance in functionality among modules is higher in better environments, rendering the probability that an advantageous mutation falls in a saturated module higher in better environments.

Our model is inspired by the modular epistasis model (Tenaillon, et al. 2012; Kryazhimskiy, et al. 2014) proposed to explain a phenomenon related to diminishing returns–a reduction in beneficial mutation rate when a population gradually rises in fitness during adaptation (Silander, et al. 2007; Tenaillon, et al. 2012). This phenomenon may be termed decreasing supplies, because it is about decreasing supplies of beneficial mutation as adaptation progresses. The modular epistasis model asserts that a population has limited ways to adapt and will run out of beneficial mutations if all modules reach their maximal functionalities. It is clear that our modular life model is similar to the modular epistasis model despite that they are proposed to explain different phenomena; one main difference is that our model includes environmental contributions to the functionalities of individual modules, allowing considering both genotype and environment qualities in the study of diminishing returns. It is also obvious that our model is able to explain decreasing supplies, because an advantageous mutation will no longer be visible to selection when its benefit reduces to a certain level via diminishing returns. This can indeed be seen in our simulation of the modular life model (**Fig. S4**).

Although the modular epistasis model is supported by the finding that deleting different genes are rescued by different sets of beneficial mutations (Blank, et al. 2014; Filteau, et al. 2015), it was disfavored in an empirical test (Kryazhimskiy, et al. 2014). Specifically, Kryazhimskiy et al. evolved *S. cerevisiae* for 240 generations to obtain 64 different founder lines. They then evolved the 64 founders for 500 generations, with 10 replicates per founder. They reasoned that, under the modular epistasis model, the substitutions observed in the 10 replicates from the same founder should have larger overlaps than those observed in the lines from different founders. However, no significant difference was detected. Although this finding appears inconsistent with the modular epistasis model, it is possible that 240 generations of evolution did not create large enough differences among the 64 founders in the distribution of functionality among modules. Another possibility is that only one module could contribute to the specific adaptation studied; therefore all improvements in all founders were in the same module, which would not predict the difference expected by the authors.

In another study (Wang, et al. 2016), several substitutions observed from an experimental evolution study of *Escherichia coli* were tested on a number of strains picked from the *E. coli* phylogeny. The authors asked whether the higher the ecological similarity between the *E. coli* strains used in the experimental evolution and tested now, the closer the growth effects of the substitutions in the two strains, but found only a marginally significant result. However, because ecological similarity may not correlate well with the similarity in module functionality, this comparison has limited power in testing the modular epistasis hypothesis.

In our simulation of the modular life model, we computed the growth rate of a genotype using either the geometric or arithmetic mean functionality among modules. While it is unclear which scenario is more appropriate, the fact that both simulation schemes qualitatively recapitulated the empirical diminishing returns patterns suggests that the primary cause of these patterns is the gene-gene and gene-environment interactions within modules. Needless to say, our simulation is oversimplified. For instance, antagonistic gene-environment interactions (Qian, et al. 2012) have not been considered. Thus, our simulation currently cannot explain how a beneficial allele becomes deleterious upon an environmental change, which is occasionally observed in real data (Wei and Zhang 2017). The modular life model is meant to provide the primary mechanism of diminishing returns. Refinement of the model with many more parameters would be necessary for it to explain the specific and detailed features of diminishing returns.

That our modular life model can recapitulate major empirical patterns of diminishing returns does not prove that it represents the truth, because the possibility exists that some other models can also explain these patterns. In this context, it is worth mentioning Fisher’s geometric model (FGM) (Fisher 1930), because it has been used to explain diminishing returns epistasis during adaptive walks (Blanquart, et al. 2014). The FGM depicts a particular, simple phenotype-fitness map without empirical basis. Under the assumption that the phenotypic effect of a mutation is independent of the genetic background, one could show that, as the background genotype becomes fitter, the benefit of a mutation reduces simply because it tends to overshoot the optimum, resulting in diminishing returns. However, mutations are highly idiosyncratic under the FGM (Tenaillon 2014), which appears inconsistent with the empirical patterns of diminishing returns (Kryazhimskiy, et al. 2014). It is also worth noting that adaptive trajectories simulated under the NK model show negative epistasis between non-consecutive substitutions and positive epistasis between consecutive substitutions (Draghi and Plotkin 2013; Greene and Crona 2014). But the prevalence of diminishing returns epistasis predicted by the NK model is much lower than observed in experimental evolution (Wünsche, et al. 2017). Whether the NK model can explain our findings from standing genetic variants in single and multiple environments is unknown.

Although our modular life model is designed retrospectively to explain patterns of diminishing returns, it can also explain several reported phenomena of mutational effects in different environments. For instance, Chou et al. tested the growth effects of a novel transporter system that enhances metal uptake in *Methylobacterium extroquens* in various metal-poor (MP) environments (Chou, et al. 2009). They observed that the same beneficial mutation had larger effects in better environments. At first glance, this observation appears contradictory to our model. However, the environments considered in our simulation of the modular life model do not have a limiting factor as in their experiment. If we consider metal uptake as a module and if the contributions of all tested MP environments to that module are equally low, our model can explain their observation. Let us assume that the product of functionalities of all modules except the metal uptake module is *M*_1_ in a relatively good environment and *M*_2_ in a relatively poor environment, respectively. Let us further assume that the environmental and genetic contributions to the functionality of the metal uptake module total x for the background genotype in all MP environments. The contribution of the beneficial mutation to the metal uptake module is *y*. Under the assumption that the growth rate is the geometric mean of all *K* modules, the growth improvement from the mutation in the relatively good environment is [*M*_1_(*x*+*y*)]^1/*K*^ - (*M*_1_*x*)^1/*K*^ = 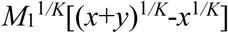. Similarly, the growth improvement from the mutation in the relatively poor environment is 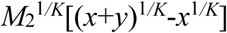. Because *M*_1_ is greater than *M*_2_, the effect size of the mutation increases as the environment gets better. The same trend is predicted by our model when the genotype instead of environment is improved in non-metal uptake modules, as was observed (Chou, et al. 2009). The phenomenon that environmental stresses sometimes decrease the harm of deleterious mutations (Kishony and Leibler 2003) can be similarly explained by our model. Note that the observations from these experiments cannot be explained if fitness equals the arithmetic mean functionality in the modular life model, suggesting that the geometric mean functionality definition of fitness may be more generally applicable than the arithmetic mean definition.

Our findings about the patterns and mechanistic basis of diminishing returns have several important evolutionary implications. First, the observation that the benefit of an advantageous mutation generally decreases with environment quality *Q* implies a negative correlation between a population’s additive genetic variance in growth rate (*V*_R_) and *Q*. This is indeed true in the yeast data (*ρ* = −0.56, *P* = 8×10^-5^; see Materials and Methods). All else being equal, the growth rate variance among individuals is also expected to decrease as *Q* rises. Consistently, we observed a negative correlation between the growth rate variance among the 1005 segregants studied here and *Q* (**Fig. 4a**). That is, the among-individual variation in growth rate gets larger as the environment becomes harsher, echoing earlier observations made in much smaller datasets (Lewontin and Matsuo 1963; Kondrashov and Houle 1994; Korona 1999; Szafraniec, et al. 2001). Second, Fisher’s Fundamental Theorem of natural selection states that the rate with which a population adapts equals the variance of fitness (Fisher 1930). Because the variance of fitness (or growth rate) rises as *Q* reduces, the same population should adapt faster in harsher environments. Third, related to the above point, evolvability is the ability of a population to respond to selection. Evolvability (*E*) equals additive fitness variance *V*_F_ divided by the mean fitness of the population (*F*) (Houle 1992). If we regard growth rate as a proxy for fitness, we have *E* ≈ *V*_R_/*Q*. Thus, evolvability rises precipitously as a population moves to harsher environments (**Fig. 4b**). This prediction is supported by some anecdotes in the literature. For instance, it was reported that the relative fitness gain in the laboratory evolution of an *E. coli* strain is faster in the less preferred temperatures of 32°C and 42°C than in its optimal temperature of 37°C (Bennett, et al. 1992). A more recent experimental evolution study also showed that the relative fitness of yeast rises more rapidly in a high temperature environment than in an optimal temperature environment (Jerison, et al. 2017). Future studies are required to test this prediction critically and systematically. Fourth, the modular structure of life creates functional redundancy within modules when the functionality of the module approaches its maximum. This redundancy means that when a population is fully adapted to an environment, the population can accumulate genetic variation with little fitness variation, a phenomenon known as phenotypic robustness to mutations (de Visser, et al. 2003; Wagner 2005). This hidden genetic variance can be useful for adaptation when the environment changes. Thus, via the phenomenon of diminishing returns, the modular structure of life fundamentally impacts both the robustness and evolvability of organisms. It will be of great interest to verify our yeast-based observations in other species.

**Fig. 4.**
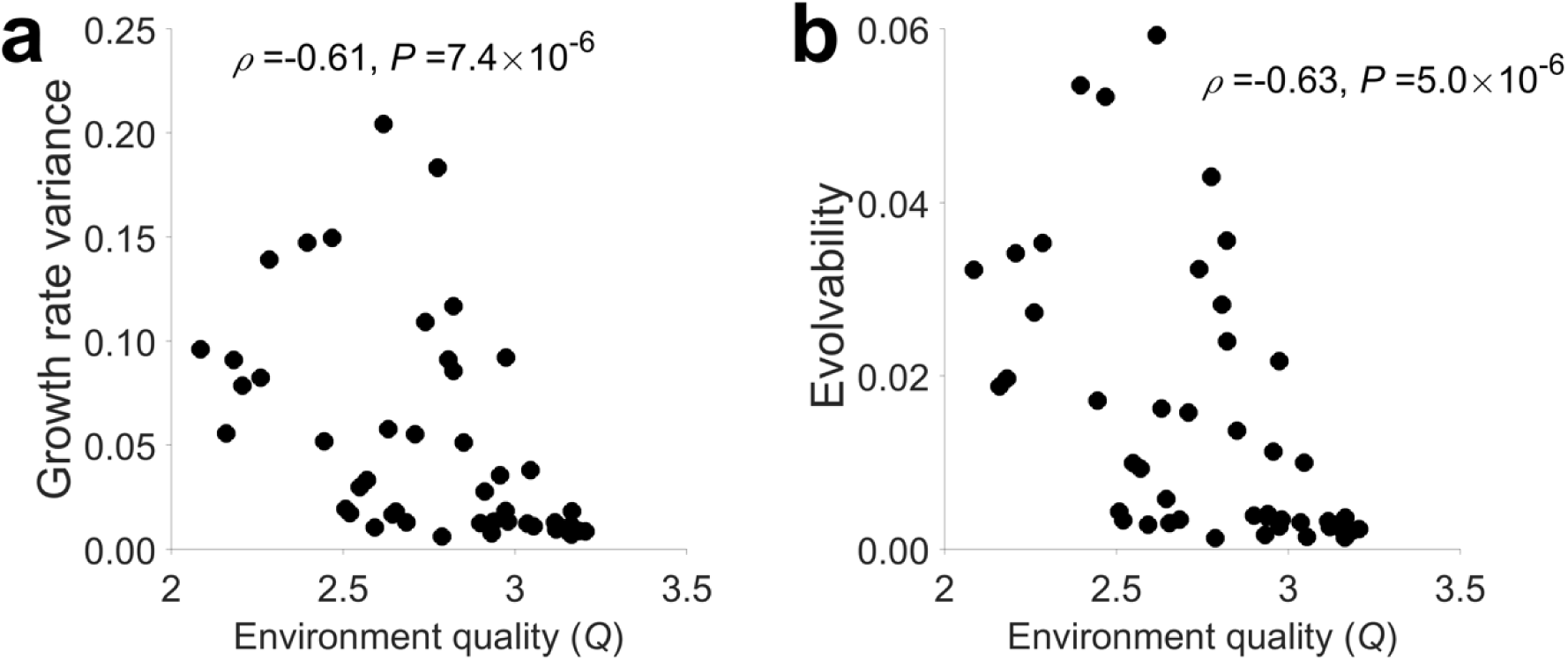
Growth rate variance and evolvability of a population increase as the environment quality (*Q*) declines. (**a**) Correlation between *Q* and the growth rate variance among the segregants examined. (**b**) Correlation between *Q* and the evolvability of the population of segregants studied. Spearman’s rank correlation *ρ* and associated *P*-value are presented.

## MATERIALS AND METHODS

### Genotype and phenotype data

We acquired from the Kruglyak lab the genotype data of 1040 segregants from a cross between the BY and RM strains of *S. cerevisiae* (Bloom, et al. 2013), including a total of 28,220 SNPs mapped to the reference genome sequence R64-1-1 (Bloom, et al. 2015). These SNPs were considered in our analysis. We downloaded the genome annotations for R64-1-1 from Ensembl biomart. We also obtained from the Kruglyak lab the average end-point colony radius of each segregant in each of 47 environments, including one (fructose medium) that was not in the original paper (Bloom, et al. 2013). After requiring each segregant to have both genotype and phenotype data in at least one environment, we retained 1005 qualified segregants for subsequent analysis. Note that colonies with ln(radius) > 3.508 had been excluded from the data by the original authors (Bloom, et al. 2013) to minimize the effect of growth saturation on growth rate estimation. We further removed those colonies with ln(radius) < 1.6, because this value approaches the lower limit of colony size measurement. To avoid potential biases created by these removals, in any environment, we considered only those SNPs for which the fraction of BY-allele-carrying segregants after the removals is not significantly different from that in all 1005 segregants (nominal *P* > 0.05, binomial test). We converted colony radius to average growth rate as described in the next section.

Growth rate variance (*V*) among segregants under each environment was computed from the growth rates of the segregants. We obtained the narrow-sense heritability (*h*^2^) under each environment from Table S2 of a previous study (Bloom, et al. 2013) and computed the additive growth rate variance by *V*_R_ = *Vh*^2^. Evolvability was calculated using *E* ≈ *Vh*^2^/*Q* (Houle 1992).

### Growth rate estimation from colony size

The original phenotype measured is the mean radius (*D*) of each colony at the end of *T* = 48h of growth on solid media. We transformed *D* to average growth rate in the following way. Let the number of cells in a colony be *N*, which can be described by

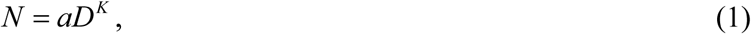

where *K* is a constant presumably between 2 (if colonies resemble columns) and 3 (if colonies resemble spheres) and *a* is a constant representing the number of cells per unit volume. Cell growth can be described by

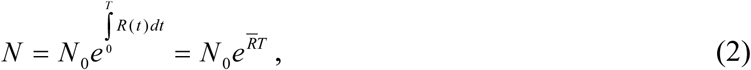

where *N*_0_ is the number of colonizing cells, *N* is the number of cells at time *T*, *R*(*t*) is the growth rate at time *t*, and *R* is the average growth rate from time 0 to *T*. From Eqs. (1) and (2), we have

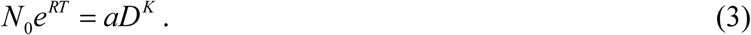

Eq. (3) can be converted to

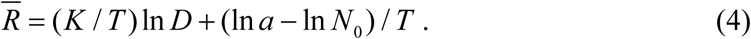

It is reasonable to assume that *N*_0_ is constant or has little effect on the colony size at 48h, because otherwise QTLs would not have been identified. Because *T* is also constant and *K* and *a* are presumably approximately constant (see below), ln*D* and *R* have approximately the same linear relationship for all strains. As a result, ln*D* can be used to represent *R* when comparing *R* values. Throughout this study, we used ln*D* as a measure of *R*. Because all analyses are based on *R*, our results hold when the average growth rate in the 48h is regarded as a fitness proxy.

To verify that *K* and *a* are approximately constant, we grew 91 randomly picked segregants on YPD agar plates for 48h. We scanned colonies and measured the pixel number per colony using SGATools (Wagih, et al. 2013), allowing quantifying the colony radius *D*. We then estimated the corresponding cell number *N* in each colony using flow cytometry (BD Accuri^TM^ C6). If *K* and *a* in Eq. (1) are constant across genotypes, ln*N* should be a linear function of ln*D*. Indeed, our data showed that ln*N* and ln*D* have a linear correlation of *r* = 0.74 (*P* < 10^-16^), supporting approximate constancies in *K* and *a* across genotypes.

To verify that the yeast growth did not saturate at 48h, we grew 79 randomly picked segregants on an YPD plate. We scanned colonies and estimated *D* at 13 time points every 2-3h from 15h to 48h of growth. The ln*D* shows a fairly linear increase over the assayed time without dramatic decline in any of the 79 segregants (**Fig. S5a**). We conducted a linear regression between ln*D* and time of growth for each colony (**Fig. S5b**), and found that the average adjusted *r*^2^ = 0.94, suggesting that *R*(*t*) did not change much during the course of 48h growth. Indeed, a quadratic fitting improves the adjusted *r*^2^ only slightly to an average of 0.96, despite that the improvement occurred to most segregants (**Fig. S5b**). Because our formulation (Eq. 4) considers the average growth rate from 0 to 48h, our method is valid as long as the slight saturation is not more pronounced for fast-growth segregants than slow-growth segregants. Indeed, we found no significant correlation among the 79 segregants tested between the growth rate rank at 48h and Δ(adjusted *r*^2^), which is the difference in adjusted *r*^2^ between the quadratic and linear regressions and a measure of saturation (**Fig. S5c**).

### Estimating epistasis from growth rate

Let *F*_WT_, *F*_A_, *F*_B_, and *F*_AB_ be the fitness of the wild-type, mutant A, mutant B, and the corresponding double mutant, respectively. It is commonly thought that (*F*_AB_/*F*_WT_) = (*F*_A_/*F*_WT_)(*F*_B_/*F*_WT_) when there is no epistasis. In other words, ln(*F*_AB_) = ln(*F*_A_) + ln(*F*_B_) - ln(*F*_WT_) under no epistasis. Let *R*_WT_, *R*_A_, *R*_B_, and *R*_AB_ be the growth rates of the wild-type, mutant A, mutant B, and the corresponding double mutant, respectively. The relationship between fitness and growth rate of a genotype is *F* = *e^Rt^*, or ln*F* = *Rt*, where *t* is the generation time of the wild-type. Hence, under no epistasis, *R*_AB_ = *R*_A_ + *R*_B_ - *R*_WT_. In other words, epistasis can be estimated by *R*_AB_ - (*R*_A_ + *R*_B_ - *R*_WT_) = (*R*_AB_ - *R*_A_) - (*R*_B_ - *R*_WT_), which is the growth effect of mutation B on the background of mutant A minus the corresponding effect on the wild-type background. This is why diminishing returns epistasis is commonly assessed by comparing the growth effect of a mutation on two genetic backgrounds.

### Assessing the fitness effect of a mutation in multiple genetic backgrounds

Diminishing returns epistasis is conventionally demonstrated by a higher growth benefit of a mutation in a less fit genotype than in a fitter genotype. Here we show that it can also be demonstrated by a higher growth benefit in a group of less fit genotypes than in a group of fitter genotypes. Suppose we are interested in assessing the growth effect of mutating allele X_1_ to X_2_ at a site in two different genetic backgrounds G and H (locus X is not considered part of the genetic background). The growth rate of the genotype with X_1_ in background G is *R*(G+X_1_) = *A*(G)+*A*(X_1_)+*E*(G)+*E*(G, X_1_), where *A*(G) is the total additive effect of all alleles in G, *A*(X_1_) is the additive effect of X_1_, *E*(G) is the total epistatic effect among all alleles in G, and *E*(G, X_1_) is the epistatic effect between X_1_ and G. Similarly, the growth rate of the genotype with X_2_ in background G is *R*(G+X_2_) = *A*(G)+*A*(X_2_)+*E*(G)+*E*(G, X_2_). Thus, the growth effect of the mutation in background G is *R*(G+X_2_)-*R*(G+X_1_) = *A*(X_2_)-*A*(X_1_)+*E*(G, X_2_)-*E*(G, X_1_) = *A*(X_2_)-*A*(X_1_)+*ΔE*(G, X_2_-X_1_), where *ΔE*(G, X_2_-X_1_) is the difference in epistatic effect between X_2_ and X_1_ in G and will be referred to as the epistatic effect of the mutation in G. The corresponding growth effect of the mutation in background H is *R*(H+X_2_)-*R*(H+X_1_) = *A*(X_2_)-*A*(X_1_)+*ΔE*(H, X_2_-X_1_). Hence, the difference between the growth effect of the mutation in H and that in G is *μ* = [*R*(H+X_2_)-*R*(H+X_1_)]-[*R*(G+X_2_)-*R*(G+X_1_)] = *ΔE*(H, X_2_-X_1_)-*ΔE*(G, X_2_-X_1_), which is the difference in the epistatic effect of the mutation in the two backgrounds. Analysis of diminishing returns is to study μ. Specifically, diminishing returns means that, when *R*(H+X_1_) > *R*(G+X_1_), *μ* = *ΔE*(H, X_2_-X_2_)-*ΔE*(G, X_2_-X_1_) < 0. In other words, when the genetic background becomes fitter, the epistatic effect of the mutation becomes smaller.

Now let us consider a group of 2*k* relatively unfit random genotypes, of which G_1_, G_2_, …, and G_k_ carry X_1_ while G_k+1_, G_k+2_, …, and G_2k_ carry X_2_; frequencies of alleles at other loci are not different between the first and last *k* genotypes. The mean growth effect of muting X_1_ to X_2_ in the above 2*k* genotypes is

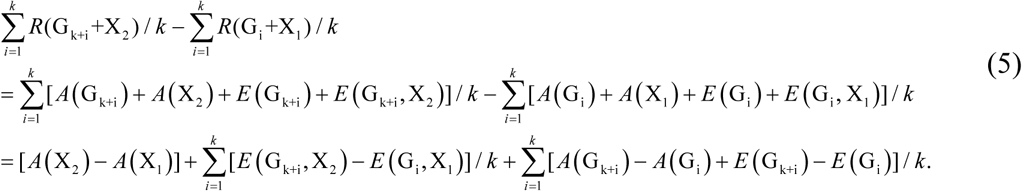

There are three terms in the right-hand side of Eq. (5). The first term is the additive effect of the mutation. The second term is the mean epistatic effect of the mutation in the genetic backgrounds concerned. The third term is expected to be 0, because the first and last *k* genotypes are on average the same in additive and epistatic growth effects. Thus, Eq. (5) can be written as

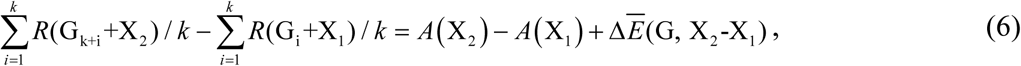

where the last term is the mean epistatic effect of the mutation in G backgrounds.

Let us similarly consider a group of 2*k* relatively fit genotypes, of which H_1_, H_2_, …, and H_k_ carry X_1_ while H_k+1_, H_k+2_, …, and H_2k_ carry X_2_. The mean growth effect of mutating X_1_ to X_2_ in the above *2k* genotypes can be similarly written as

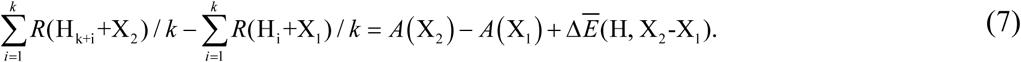

Using Eqs. (6) and (7), we can find that the difference between the growth effect of the mutation in the H backgrounds and that in the G backgrounds is

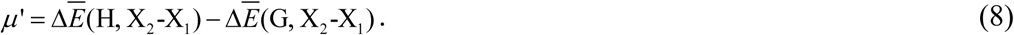

Thus, it is clear that *μ* and *μ’* measure the same thing except that the epistatic effect of the mutation in one genetic background is considered in the former while the mean epistatic effect of the mutation in multiple backgrounds is considered in the latter. Given the stochasticity of epistasis, mean epistasis is presumably more informative than a single epistasis value in the study of diminishing returns patterns.

### Bootstrap test of the significance of diminishing returns epistasis

We examined whether *s*_H_ is significantly smaller than *s*_L_ for each QTL by a bootstrap test. We first calculated the observed *s*_L_ - *s*_H_. We then generated a bootstrap sample of growth rates from the fitted 20% of BY-carrying segregants as well as a bootstrap sample of growth rates from the fitted 20% of RM-carrying segregants, allowing the estimation of *s*_H_ from the bootstrap samples. We similarly generated bootstrap samples and obtained the estimate of *s*_L_ and then *S*_L_-*s*_H_. This process was repeated 10,000 times. *P*-value is estimated by the proportion of bootstrap replications in which *s*_L_ < *s*_H_.

### Testing the effect of SNP density on *Q-s* correlation

For every 20 consecutive SNPs, we calculated the distance in base pair between the two boundary SNPs and then the SNP density in the region. We calculated the fraction of SNPs showing negative *Q-S* correlations as well as the average correlation coefficient of these 20 SNPs. Among 1396 regions with 20 SNPs, SNP density does not correlate with the fraction of SNPs showing negative *Q-s* correlations in the region (*ρ*= 0.0161, *P* = 0.55), nor the average correlation coefficient of the 20 SNPs (*ρ*= −0.045, *P* = 0.12). Similar results were obtained when 549 regions each with 50 consecutive SNPs were analyzed (*ρ*= 0.045 and −0.050; *P* = 0.29 and 0.24, respectively).

### Simulation of the modular life model

We assume that the growth rate of a genotype in an environment is the combined effects of *C* functional modules. Each module has a functionality value that is the sum of environmental and genetic contributions to the module. The maximum possible functionality of each module is 1 and the minimum is 0. Consequently, further improvement in genotype or environment quality has no contribution to the functionality of a module when it reaches the maximum. Each module has *M* contributing genes, each with one SNP that distinguishes between a fully functional allele and a null allele. There are *N* haploid segregants in a population; the genotype of each segregant is made up of *CM* genes, each carrying the functional allele with a 50% probability.

In our simulation, the specific values of various parameters are not critical to the conclusion, as long as the functionalities of some modules reach the upper limit. Below is the set of parameters used in generating Fig. 3bc. We used *C* = 10, *M* = 10, and *N* = 1000, and simulated 50 environments. The maximal contributions of the 10 genes to the functionality of a module were set to be 0.11, 0.12, 0.13, …, and 0.2, respectively. Thus, the functional allele of gene 1 contributes 0.11 units of functionality to its module, while the null allele contributes 0 unit. We assumed that the contribution of an environment to a module is a normal random variable with a standard deviation of 0.05. The mean of the normal distribution is 0.2000, 0.2035, 0.2070, …, and 0.3715, respectively, from the 50 environments. We also added a noise
term, drawn randomly from the normal distribution of mean = 0 and standard deviation = 0.01, to the growth rate of each simulated genotype in each environment. The simulation parameters when fitness equals the arithmetic mean functionality across modules are detailed in **Fig. S2** legend.

### Reanalysis of Kryazhimskiy et al.’s data of diminishing returns

We reanalyzed the data from Figure 3 of Kryazhimskiy et al. (Kryazhimskiy, et al. 2014). The growth rates of all strains were measured using flow cytometry-based competition assays against the ymCitrine-labelled DivAncCit strain and were represented by percent difference from DivAncCit. *HO, GAT2, WHI2*, and *SFL1* genes were separately deleted in each of 40 different ancestor strains. The growth rate of each ancestor strain was measured in triplets, and we calculated the mean growth rate and its standard error using the three repeats. For the deletion strains, the growth rates of one to five replicate colonies were measured three times each. For these strains, we first calculated the growth rate of each replicate and then calculated the mean growth rate and its standard error using the replicates. When there was no replication, we calculated the mean growth rate and its standard error using the repeats. We used two-tailed *Z*-test to identify all pairs of strains whose growth rates are not significantly different from each other. For each of these strain pairs, we used a two-tailed *Z*-test to test if the effect sizes of the same mutation are significantly different. The strain pairs with significantly different growth effects for the same mutation after Bonferroni correction are shown in **Table S2**.

### Data availability

All data generated in this study and scripts for modular life model simulation are available under https://github.com/AprilWei001/diminishing-Returns-Epistasis.

## ACKNOWLEDGEMENTS

We thank the Kruglyak lab for sharing the yeast segregant genotype and phenotype data and Alexey Kondrashov and members of the Zhang lab for valuable comments on earlier versions. This work was supported in part by the U.S. National Institutes of Health grant R01GM120093 to J.Z.

